# IL-6 signalling protects zebrafish larvae during *Staphylococcus epidermidis* infection in a novel bath immersion model

**DOI:** 10.1101/2020.02.26.960443

**Authors:** PT Dhanagovind, Prabeer K. Kujur, Rajeeb K. Swain, Sanjita Banerjee

**Affiliations:** School of Biological Sciences, National Institute of Science Education and Research, Bhubaneswar; Institute of Life Sciences, Bhubaneswar

**Keywords:** *Staphylococcus epidermidis*, zebrafish, miRNA-142, IL-6, innate immune signalling

## Abstract

Host immune responses to *Staphylococcus epidermidis*, a frequent cause of nosocomial infections, are not well understood. We have established a novel bath immersion model of this infection in zebrafish larvae. *S.epidermidis* infection activates Tlr-2 signalling pathway by upregulation of *tlr-2*. Macrophages play a primary role in the host immune response and are involved in clearance of infection in the larvae. There is marked inflammation characterised by heightened NF-κB signalling and elevation of several pro-inflammatory cytokines. Infected larvae show rapid upregulation of *il-1b* and *tnf-a* transcripts and relatively slower elevation of *il-6* transcription. The IL-6 signalling pathway is additionally subject to amplification by elevation of IL-6 signal transducer (*il-6st*) levels, which negatively correlates with miRNA dre-miR-142-5p expression. Enhanced IL-6 signalling is protective to the host in this model as inhibition of the signalling pathway resulted in increased mortality upon *S.epidermidis* infection. Our study describes the host immune responses to *S.epidermidis* infection, identifies a likely role for miR-142-5p – *il-6st* interaction in modulating this response and establishes the importance of IL-6 signalling in this infection model.

## Introduction

Initially thought to be innocuous commensals, *Staphylococci* are increasingly being recognised as pathogens that are responsible for the aetiology of a number of opportunistic human and animal diseases (Chessa et al., 2015). Classified as either coagulase-positive staphylococci (CoPS) or coagulase-negative staphylococci (CoNS) based on presence or absence of enzyme coagulase, *S.aureus* and *S.epidermidis* infections rank at the top of causative agents for nosocomial infections (Chessa et al., 2015). *Staphylococci* have also been observed to rapidly gain antibiotic resistance (Marsilio et al., 2018). Hence it is important to understand the virulence mechanisms and to decipher the host immune responses.

Research on *Staphylococci* has mostly focussed on *S.aureus*, which is a leading cause of bacteraemia worldwide. *S.aureus* is a CoPS bacterium with an extensive arsenal of virulence factors (Gordon and Lowy, 2008). *S.epidermidis*, a CoNS bacterium, has remained relatively less-studied, plausibly due to the fact that *S.epidermidis* infection is usually not life-threatening. Although *S.epidermidis* lacks aggressive virulence determinants, the genome possesses a wide range of genes that equip the organism to survive in harsh conditions in order to establish a commensal relation with its host (Otto, 2009). This gives the bacterium an added advantage during infections (Otto, 2009). *S.epidermidis* shows high diversity, with 74 identified sequence types. Ironically, there is no genomic data available for the isolate that has most commonly been isolated and is thought to be potentially the most invasive (Otto, 2009). Multi-drug resistant strains of *S.epidermidis* are quite widespread. Nearly 76% of clinical samples obtained from patients with catheter-related bacteraemia were multi-drug resistant (Cherifi et al., 2013). While the prevalence of drug resistance is low in North America, it is as high as 40% in Asian and African populations (Morgenstern et al., 2016; Sabaté Brescó, Marina et al., 2017).

*S.epidermidis* is the most frequent pathogen causing infections in indwelling medical devices as well as early-onset neonatal sepsis (Otto, 2009; Widerström, 2016). The ability of this bacterium to form biofilm further protects it from host immunity as well as antimicrobial treatment regimens (Le et al., 2018). *S.epidermidis* infections often lead to chronic infections, and occasionally to acute sepsis; the latter especially in neonates (Cheung and Otto, 2010; Nguyen et al., 2017). The high prevalence of multi-drug resistant strains, particularly in the clinical setting as well as the increasing incidence of such infections have made *S.epidermidis* infections a serious burden for health-care world-wide. For example, *S.epidermidis* alone accounts for ~22% of bloodstream infections in intensive care setting in the US, ~30% of orthopaedic-device related infections that may increase up to ~50% in developing countries, and 15-40% of prosthetic valve endocarditis (Le et al., 2018; Otto, 2009; Sabaté Brescó, Marina et al., 2017).

Mechanisms of *S.epidermidis* biofilm formation and immune responses to the infection have been investigated in cell cultures and ex vivo models. Characterisation of host immune responses to this pathogen in vivo however has been scarce (Sabaté Brescó, Marina et al., 2017). Here we report the establishment of a highly amenable in vivo zebrafish model of *S.epidermidis* infection. We demonstrate an early cellular response that is composed of macrophages and a subsequent immune response that upregulates the Tlr-2 signalling pathway. The host mounts an NF-κB mediated pro-inflammatory response with an upregulation of the IL-6 signalling pathway that has an important protective role in host immunity. Further we identify a miRNA mechanism that potentially has a role in IL-6 signal amplification during this infection.

## Results

### Establishment of a bath immersion model of *S. epidermidis* infection

In order to ascertain the range of bacterial doses to which zebrafish are susceptible and the infection model is amenable to experimentation, 4 days post-fertilisation (dpf) larvae were exposed to three doses; 5 × 10^8^, 1 × 10^9^, 1 × 10^10^ CFU/ ml of *S.epidermidis* for 6- and/ or 24 h and survival was monitored till 80 hours post-infection (hpi) (**Fig - 1A**). The larvae incubated with the highest dose of bacteria for 6-hours (Inf–high-6h) showed severe mortality (inverted triangles, open). As early as 8 hpi, survival dropped to 65% (**, p = 0.002), and by 16 hpi less than 48% survived (****, p < 0.0001). For the doses 5 × 10^8^ CFU/ ml (squares, open; Inf–low-6h) and 1 × 10^9^ CFU/ml (circles, open; Inf–med-6h), a 6-hour exposure resulted in 95% and 83% larvae surviving respectively. These mortality values were not significantly different with respect to the control population. However, when the exposure window was increased to 24 hours, both these infection doses (Inf-low-24h and Inf-med-24h) caused significant mortalities. At 16 hpi, larvae exposed to Inf-med-24h (squares, closed) had 75% survivors (*, p = 0.048). The death steadily increased till 48 hpi when there were only 40% survivors (****, p < 0.0001). Thereafter, no further significant mortality was observed. For larvae exposed to Inf-low-24h (circles, closed), larval death was apparent from 12 hpi, but the mortality reached values significantly higher than controls at 48 hpi (*, p = 0.02). Thereafter it stabilised and at the end of the study, at 80 hpi, 73% of the larvae had survived (**, p = 0.004). In both the cases, there was minimal death observed beyond 48 hpi. Heat-killed bacteria (5 × 10^8^ CFU/ ml – grey squares; heat-killed–low-24h and 1 × 10^9^ CFU/ ml – grey circles; heat-killed–med-24h) did not cause any mortality, indicating that mortalities observed were due to *S.epidermidis* infection. These results demonstrated that within the range of dose and exposure that we have tested mortality and hence severity of the disease was dependent on the dose and duration of exposure to the bacteria. We also ascertained the range of bacterial dose as well as exposure periods over which the phenotype was consistently measurable. Given the high lethality that was observed, Inf–high-6h was not used for further studies.

**Figure – 1:**
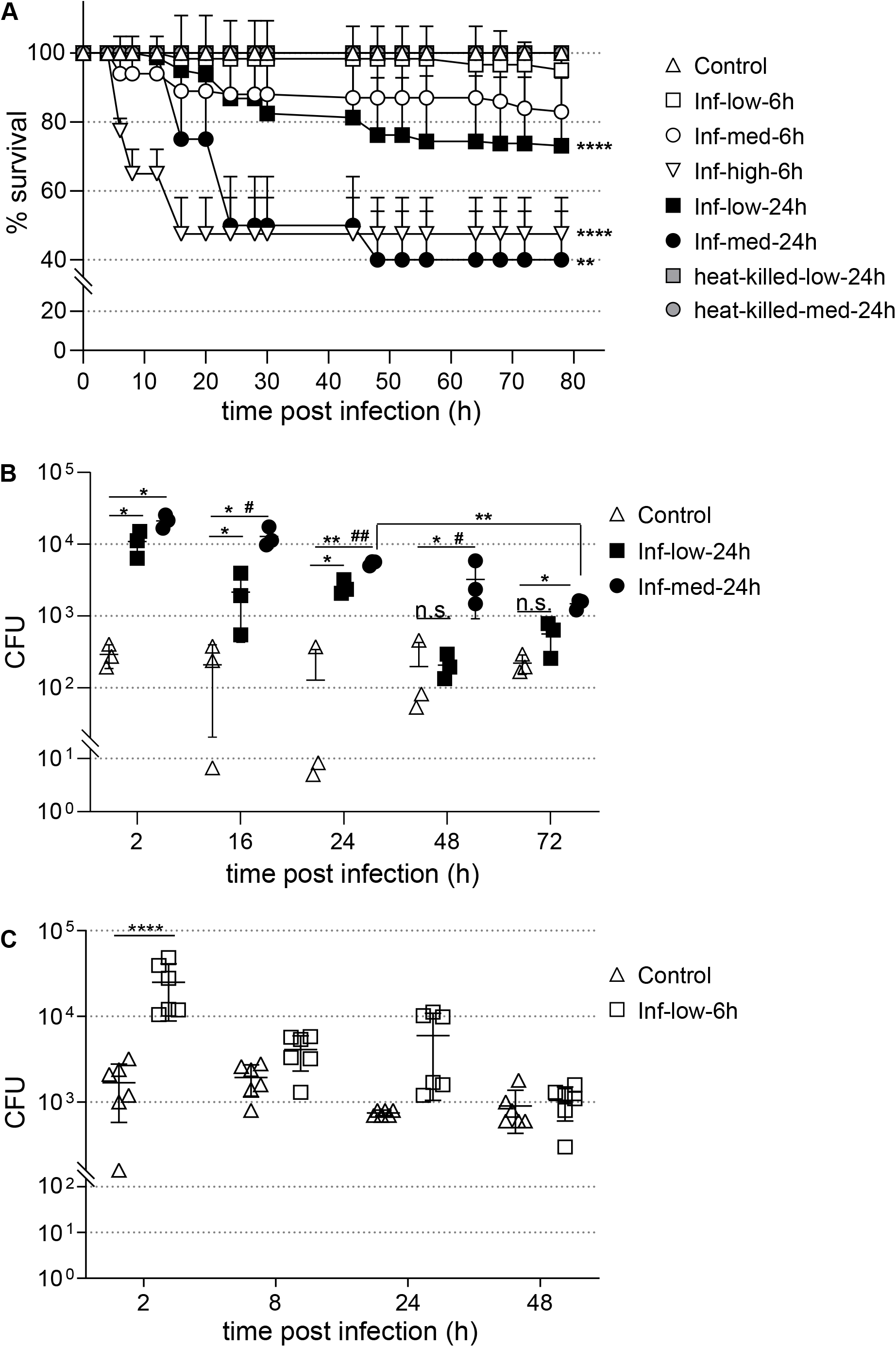
Zebrafish larvae infected with *S.epidermidis* show bacterial colonisation followed by clearance. (**A**) Larvae, 4 dpf, were exposed to varying doses (low − 5 × 10^8^ CFU/ml, med - 1 × 10^9^ CFU/ml and high - 1 × 10^10^ CFU/ml) of *S.epidermidis* for 6- or 24 h and mortality monitored. Each well had 15 larvae. Larvae exposed to low or medium doses of bacteria for 6 h (Inf-low-6h and Inf-med-6h) did not show significant mortality. Larvae exposed to high dose for 6 hours (Inf-high-6h) showed significant mortality 8 hpi onward. Longer (24 hours) exposure of medium and low doses (Inf-med-24h and Inf-low-24h) also resulted in significantly higher deaths 48 hpi and 24 hpi onward respectively (**, p < 0.01; ****, p < 0.0001). Heat-killed bacteria had no effect on zebrafish survival. (**B**) *S.epidermidis* colonisation and clearance was monitored by CFU plating of uninfected, Inf-low-24h and Inf-med-24h larvae. Compared to control larvae, the CFU counts of infected larvae remained significantly high till 24 hpi (*, p < 0.05; **, p < 0.01). Inf-low-24h showed clearance post-24 hpi, but CFU counts in Inf-med-24h larvae remained significantly elevated till 72 hpi (*, p < 0.05). Inf-low-24h had significantly lower CFU counts than Inf-med-24h between 16 to 48 hpi (#, p < 0.05; ##, p < 0.01). The CFU counts of Inf-med-24h at the at 72 hpi was significantly lower than that at 24 hpi (**, p < 0.01). (**C**) Bacterial clearance was monitored by infecting 4 dpf larvae with Inf-low–6h. Infected larvae had significantly higher CFU counts at 2 hpi (****, p < 0.0001) compared to controls. At later time-points there was no difference in the CFU counts between uninfected and Inf-low-6h larvae. Representative of three independent experiments.

### Infection dose determines bacterial clearance

We then followed the infection progress and looked at initial colonisation of *S.epidermidis* followed by its clearance over time (**Fig - 1B**). Since larvae exposed to the low and medium doses of bacteria for 6-hours did not show significant mortality, the longer exposure period (24 h) was used for this purpose. CFUs were counted at different time-points till 72 hpi from uninfected controls, Inf–low-24h and Inf–med-24h larvae. The CFU counts from the uninfected larvae remained low throughout the experiment. The infected larvae of both the doses showed significantly higher CFU counts than controls from 2 hpi till 24 hpi (*, p = 0.013 for Inf-low-24h, *, p = 0.047 at 2- and 16 hpi, **, p = 0.008 at 24 hpi for Inf-med-24h). At 2 hpi, there was no difference between the two infected groups indicating that the initial colonisation efficiency was not influenced by the dose. From the next time-point that we checked, 16 hpi onward, while both the populations showed progressively decreasing CFU yields, the larvae infected with Inf-med-24h consistently showed significantly higher CFU counts than those infected with Inf-low-24h (#, p = 0.047, ##, p = 0.0055). At 48 hpi, the CFU counts of Inf–low-24h larvae had decreased to values similar to the control population. The CFU counts of Inf–med-24h larvae however remained significantly elevated during the remainder period of the experiment, till 72 hpi (*, p = 0.011).

While the CFU counts decreased over time after exposure to Inf–low-24h, the CFU counts remained elevated and bacteria were not completely cleared till 72 hpi when exposed to Inf–med-24h. The reduction in CFU counts coincides with significant larval death. Hence it can be argued that the reduction in CFU is due to the severely infected larvae dying out leaving behind survivors with lower levels of infection, and thus is not indicative of bacterial clearance per se. This argument, however, does not explain the difference in the CFU counts that are observed between the two infection doses. The larvae infected with low dose showed lower mortalities but were able to clear bacteria faster than the larvae infected with medium dose, as evidenced by the CFU counts over time. Additionally, larvae of Inf-med-24h show reduction of bacterial counts post 24 hpi without any significant accompanying death. This can be seen in the significant decrease in CFU counts between 24 hpi and 72 hpi of this population (CFU counts: **, p = 0.0017; larval mortality: ns, p = 0.87). This strongly suggests that the zebrafish are able to clear the bacteria after the initial colonisation phase at these multiplicities of infection (MOI), and this phenomenon contributes, at least in part, to the reduction in bacterial counts in the surviving larvae. To investigate further, we tested bacterial colonisation and clearance in larvae infected with In-low-6h – a dose that showed minimal mortality and was indistinguishable from the uninfected controls (**Fig - 1C**). At 2 hpi, Inf–low-6h larvae yielded significantly higher CFU counts than their uninfected counterparts (****, p = 5.17 × 10^−8^) indicating that infection does occur at this dose. By 8 hpi, the CFU counts from the infected population had fallen to values similar to the uninfected controls. This demonstrates that zebrafish are able to clear the infecting bacteria by mounting an immune response against the infectious agent. Exposure to higher density or duration of the pathogen has deleterious effects likely by overwhelming host clearance mechanisms, and thus leading to colonisation.

### Macrophage response is induced as early as two hours post *S.epidermidis* infection

We next aimed to identify the cellular lineage(s) that might be involved in mediating the immune response to *S.epidermidis* infection. Both macrophages and neutrophils have been shown to be involved in host immune responses to Staphylococcal infections (Le et al., 2018; Rigby and DeLeo, 2012; Verdrengh and Tarkowski, 2000). Given that the leukocytic population of a 4 dpf larvae is primarily constituted of macrophages and neutrophils of myeloid lineage, we used *Tg*(*pu.1*:*RFP*), a transgenic reporter fish line to investigate the role of circulating leukocytes in the response to *S.epidermidis* infection. The expression of TagRFP in these fish is driven by the *pu.1* promoter, a myeloid-lineage specific transcription factor (Rhodes et al., 2005). Hence, the number of *pu.1*-positive cells is indicative of the total number of leukocytes in the larvae. To determine whether or not *S.epidermidis* infection resulted in a change in the total number of leukocytes, *Tg*(*pu.1*:*RFP*) larvae were infected with the Inf-med-24h dose and uninfected and infected larvae were imaged at different time-points post-infection. Higher number of *pu.1*-positive cells were seen in the infected larvae compared to the controls. In order to quantitate this, the total number of *pu.1*-positive cells in the same segment of multiple infected and control larvae were counted and the average values compared. The infected larvae showed a significant increase in the number of *pu.1*-positive cells as early as 2 hpi (**Fig - 2A-B)**. The infected larvae had an average of 24.6 ± 7.0 *pu.1*-positive cells at this time-point compared to controls (11.6 ± 5.1) (***, p = 0.0003) (**Fig - 2E**). The leukocyte numbers in the infected population remained elevated even at 24 hpi, showing a nearly 3-fold increase over the uninfected control population (**Fig - 2F**). At 24 hpi, the average number of *pu.1*-positive cells in Inf–med-24h larvae was 19.3 ± 4.5 as opposed to 6.3 ± 4.0 in controls (***, p = 0.0008) (**Fig - 2C-D, E**). While the data presented here reflect the number of *pu.1*-positive cells in the trunk region (as indicated in **Fig - 2**), similar trend was also observed in the tail region.

**Figure – 2:**
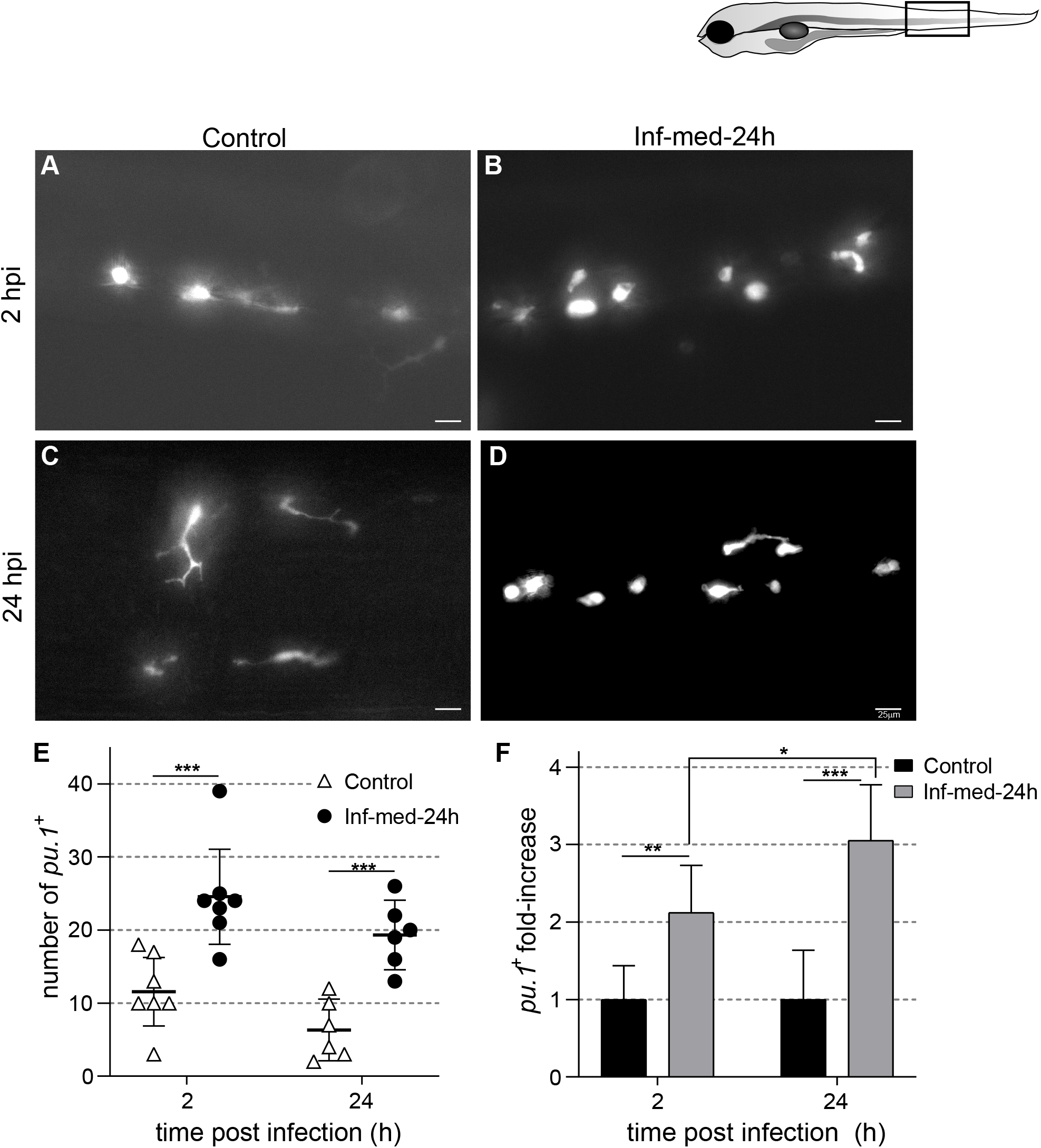
*S.epidermidis* infection results in an increase in leukocytes. *Tg* (*pu.1*:RFP), 4 dpf, were infected with Inf-med-24h dose, and number of *pu.1*-positive cells counted in similar segments of control (**A**& **C**) and infected (**B** & **D**) larvae. Significant increase in *pu.1* population was seen in infected larvae as early as 2 hpi (**B**). The population of *pu.1* cells remained high till 24 hpi (**D**). (**E**) The average number of *pu.1* cells in Inf-med-24h and uninfected controls were plotted at 2 hpi and 24 hpi (***, p < 0.001) (n = 6 - 7 larvae). (**F**) The Inf-med-24h larvae showed significant fold-increase at 2- and 24 hpi (*, p < 0.05**, p < 0.01; ***, p < 0.001). Box represents the area imaged. Images are representative of three independent experiments. Quantification reflects combined data of three independent experiments. Scale bar, 25 μm.

If leukocytosis plays a critical role in limiting and clearing the bacterial infection, then the larvae infected with Inf-low-6h, the dose that clears *S.epidermidis* asymptomatically, should also exhibit elevation in the number of *pu.1*-positive cells. Inf-low-6h larvae show an initial increase in the number of *pu.1*-positive cells followed by a gradual decrease (**Fig - 3**). At 2 hpi, more than 2-fold increase in the number of *pu.1*-positive cells was seen in Inf-low-6h larvae (22.6± 14) compared to uninfected controls (9.9 ± 5.4) (**, p = 0.0012) (**Fig - 3A-B, I, J**). This was comparable to the increase observed in the Inf-med-24h case. At 6 hpi, the infected larvae still showed 1.9-fold increase (21.8 ± 11.7 *pu.1*-positive cells) (**, p = 0.0069) (**Fig - 3C-D, I, J**), which fell to 1.7-fold by 18 hpi (20.6 ±7.9 *pu.1*-positive cells) (*, p = 0.015) (**Fig - 3E-F, I, J**). The number of *pu.1*-positive cells in uninfected larvae at these time-points was 11.0 ± 4.6 and 11.7 ± 3.7 respectively. By 24 hpi, the number of *pu.1*-positive cells in the infected larvae (13.2 ± 4.0) was similar to the uninfected larvae (12.4 ± 3.6) (**Fig - 3G-H, I, J**). The data suggested that leukocytosis is a critical component of the immune response mounted by infected larvae in our model. *S.epidermidis* infection-induced increase in circulating leukocytes appears to be concomitant with bacterial colonisation. The resolution phase of leukocytosis however tails off in time beyond the infection clearance window.

**Figure – 3:**
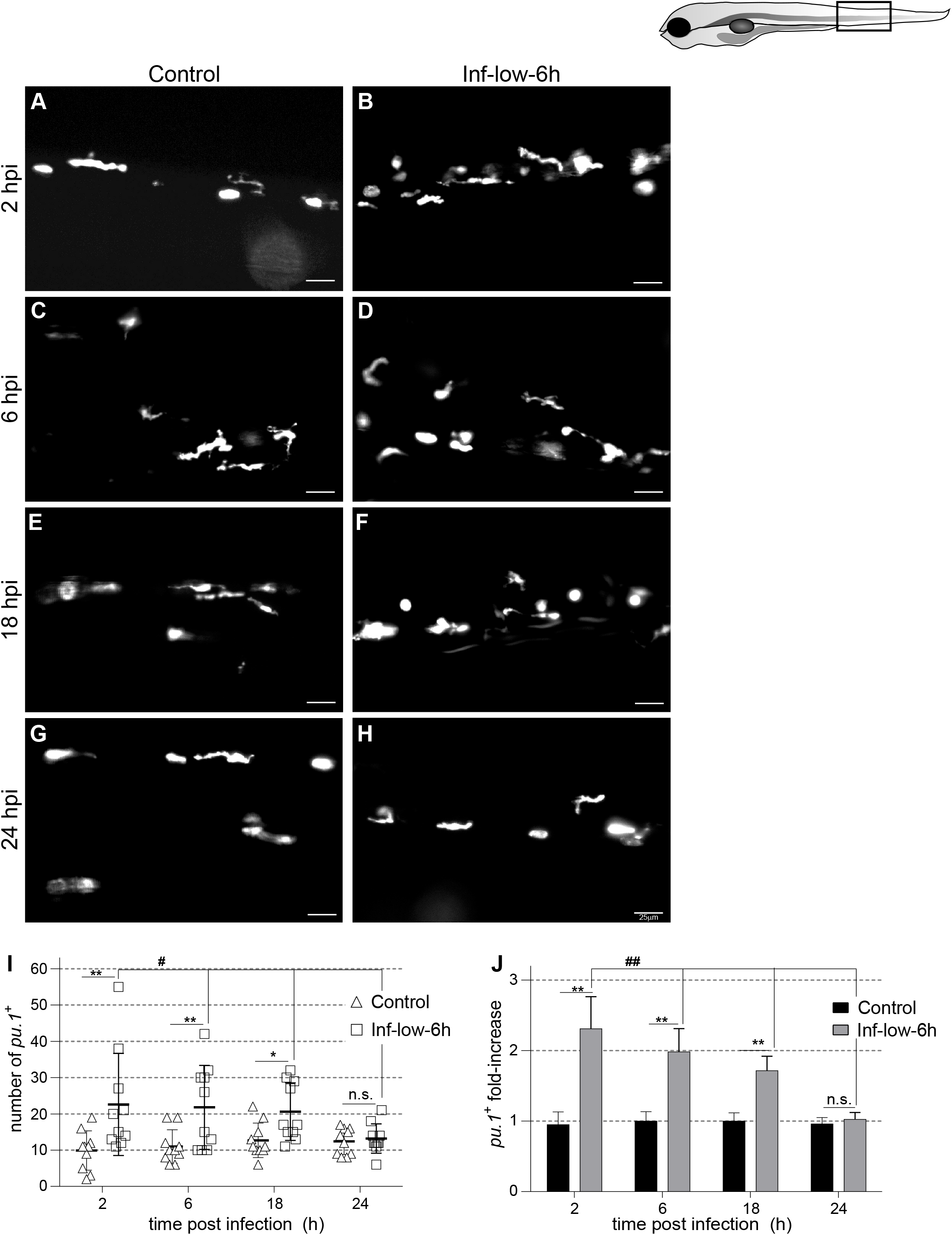
Leukocytes are critical in mediating bacterial clearance. *Tg* (*pu.1*:RFP), 4 dpf, were infected with Inf-low–6h, and number of *pu.1*-positive cells counted in similar segments of control and infected larvae at 2 hpi (**A-B**), 6 hpi (**C-D**), 18 hpi (**E-F**) and 24 hpi (**G-H**) respectively. Inf-low-6h larvae had significantly higher number of *pu.1*-cells till 18 hpi. (**I**) The average number of *pu.1* cells in Inf-low-6h and uninfected controls were plotted at each time-point (*, p < 0.05; **, p < 0.01). Compared to 24 hpi, Inf-low-6h larvae showed significant fold-increase in *pu.1*-cells at 2 hpi, 6 hpi and 18 hpi (#, p < 0.05). (**J**) The fold-increase in *pu.1*-positive cells were significant till 18 hpi (**, p < 0.01). Compared to 24 hpi, Inf-low-6h larvae showed significant fold-increase in *pu.1*-cells at 2 hpi, 6 hpi and 18 hpi (##, p < 0.01). Box represents the area imaged. Images are representative of three independent experiments. Quantification reflects combined data of three independent experiments. Scale bar, 25 μm.

Given that both macrophages and neutrophils have been implicated in host immune responses to *S.epidermidis* (Le et al., 2018), we next sought to analyse the relative contributions of the two populations to the observed leukocytosis in our model. The *pu.1*-expressing responder cell population could be macrophages, or neutrophils, or both. In order to differentiate between the two lineages, larvae obtained from mating *Tg*(*pu.1*:*RFP*) with *Tg*(*mpeg1*:*Kaede*) were used for infection and imaging experiments. Only macrophages are labelled by *Tg*(*mpeg1*:*Kaede*) (Ellett et al., 2011). These larvae were imaged under conditions where Kaede behaves as a green fluorescence protein. As can be seen from **Fig - 4**, the neutrophil population remained unaffected by *S.epidermidis* infection (**Fig - 4E**), while there was a significant increase in the number of macrophages at both 2 hpi (**Fig - 4A-B’’, E**) and 24 hpi (****, p < 0.0001) (**Fig - 4C-D’’, E**). While the average number of macrophages in Inf– med-24h larvae at 2- and 24 hpi were 22.4 ± 7.7 and 23.6 ± 3.0 respectively, the macrophage numbers in controls at the corresponding time-points were 11.9 ± 2.8 and 13.2 ± 3.6 respectively. The increase in leukocyte numbers thus mostly owes to increased number of macrophages in circulation. Hence it is likely that macrophages are the predominant cellular responders to *S.epidermidis* infection.

**Figure – 4:**
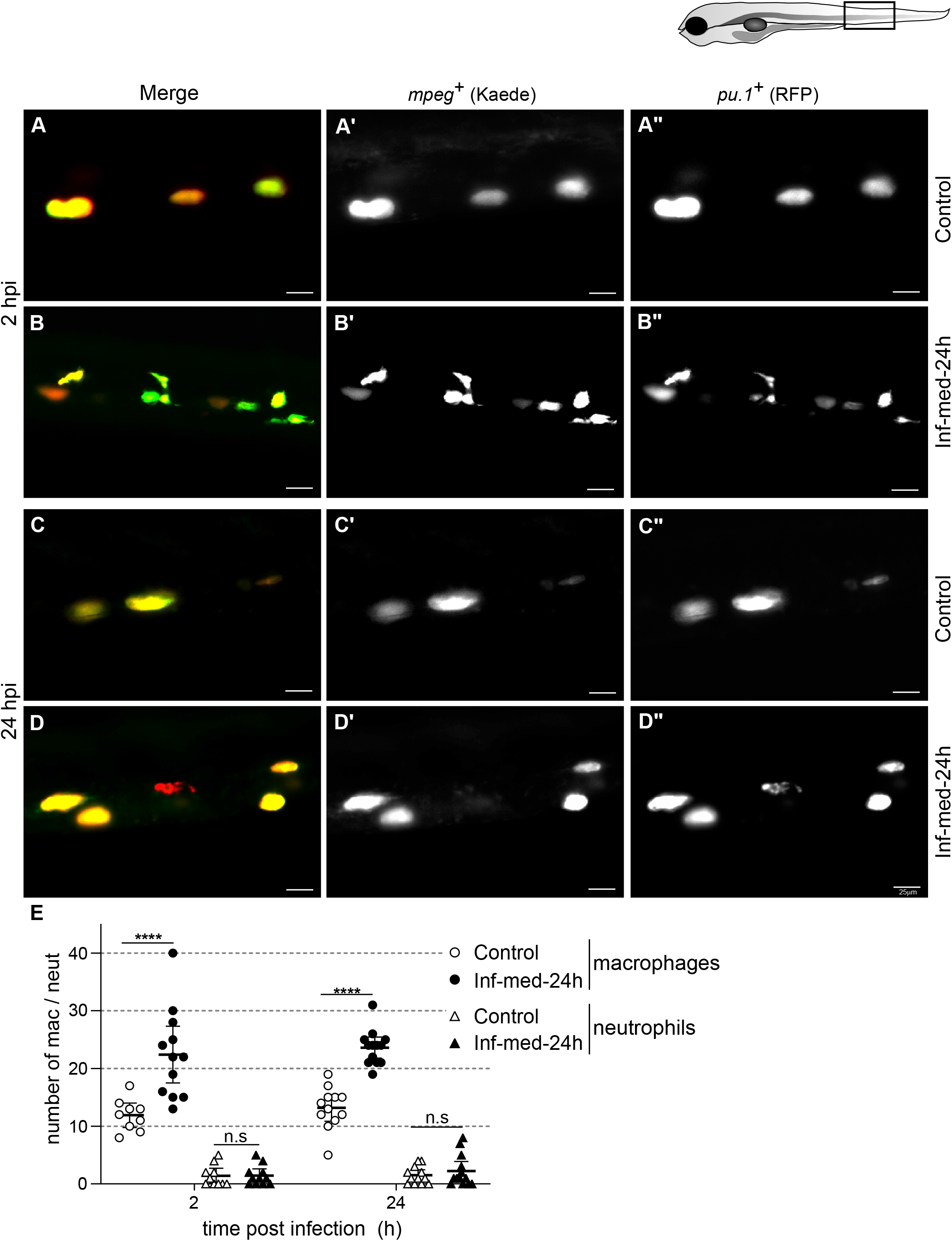
Macrophages are central responders to *S.epidermidis* infection. Double positive larvae obtained from *Tg*(*pu.1*:*RFP*) x *Tg*(*mpeg*:*Kaede*) mating were infected with Inf-med-24h and number of macrophages and neutrophils enumerated at 2- and 24 hpi at similar segments of controls (**A**-**A’’** & **C**-**C”**) and infected cohorts (**B**-**B’’**& **D**-**D”**). There was a significant increase in the macrophage population in the infected population at both 2-(**A’** vs **B’**) and 24 hpi (**C’** vs **D’**). (**E,** open and closed circles) The average number of macrophages at 2 hpi and 24 hpi were significantly elevated in infected larvae in comparison to controls (****, p < 0.0001). The neutrophil numbers were comparable between the two experimental groups, at both the time-points (**A”** vs **B’’** at 2 hpi; **C’’** vs **D’’** at 24 hpi; **E**, open & closed triangles) (n = 9 - 13 larvae). Box represents the area imaged. Images are representative of three independent experiments. Quantification reflects combined data of three independent experiments. Scale bar, 25 μm.

### *tlr-2* upregulation and NF-κB activation drive the host response to *S.epidermidis* infection

We were interested in understanding the signalling pathways that control the host response to *S. epidermidis* infection. TLRs act at the forefront of innate host defences, and *tlr-2* has been implicated in driving host immune response to Gram-positive bacteria (Takeuchi et al., 1999). Real-time qPCR experiments showed that *tlr-2* mRNA levels were significantly upregulated upon infection in our model (**Fig - 5A**). As early as 2 hpi, the Inf–med-24h larvae, compared to controls, had over 1.5-fold increase in *tlr-2* mRNA levels (**, p = 0.007). The levels increased further at 16 hpi (over 5.5-fold, ***, p = 0.0007) and remained high even at 24 hpi (over 3-fold, **, p = 0.0013). Apart from Tlr-2, involvement of Tlr-4 and Tlr-8 have been reported in murine and human cell-lines respectively in response to *S.aureus* infection (Bergstrøm et al., 2015; Stenzel et al., 2008). When we looked at *tlr-4b* levels, we noticed a modest but steady rise of this transcript in infected larvae over the uninfected controls (**Fig - 5B**). At 2 hpi, there was over 1.5-fold increase in *tlr-4b* levels. The *tlr-4b* levels rose to 2.5-fold at 16 hpi and remained elevated till 24 hpi (2.9-fold, ***, p = 0.001). The levels of *tlr-8a* and *tlr-8b* remained unaffected (**Fig S1A-B**). Thus in this infection model, pathogen recognition primarily occurs likely via *tlr-2*, with a limited role for *tlr-4b*.

**Figure – 5:**
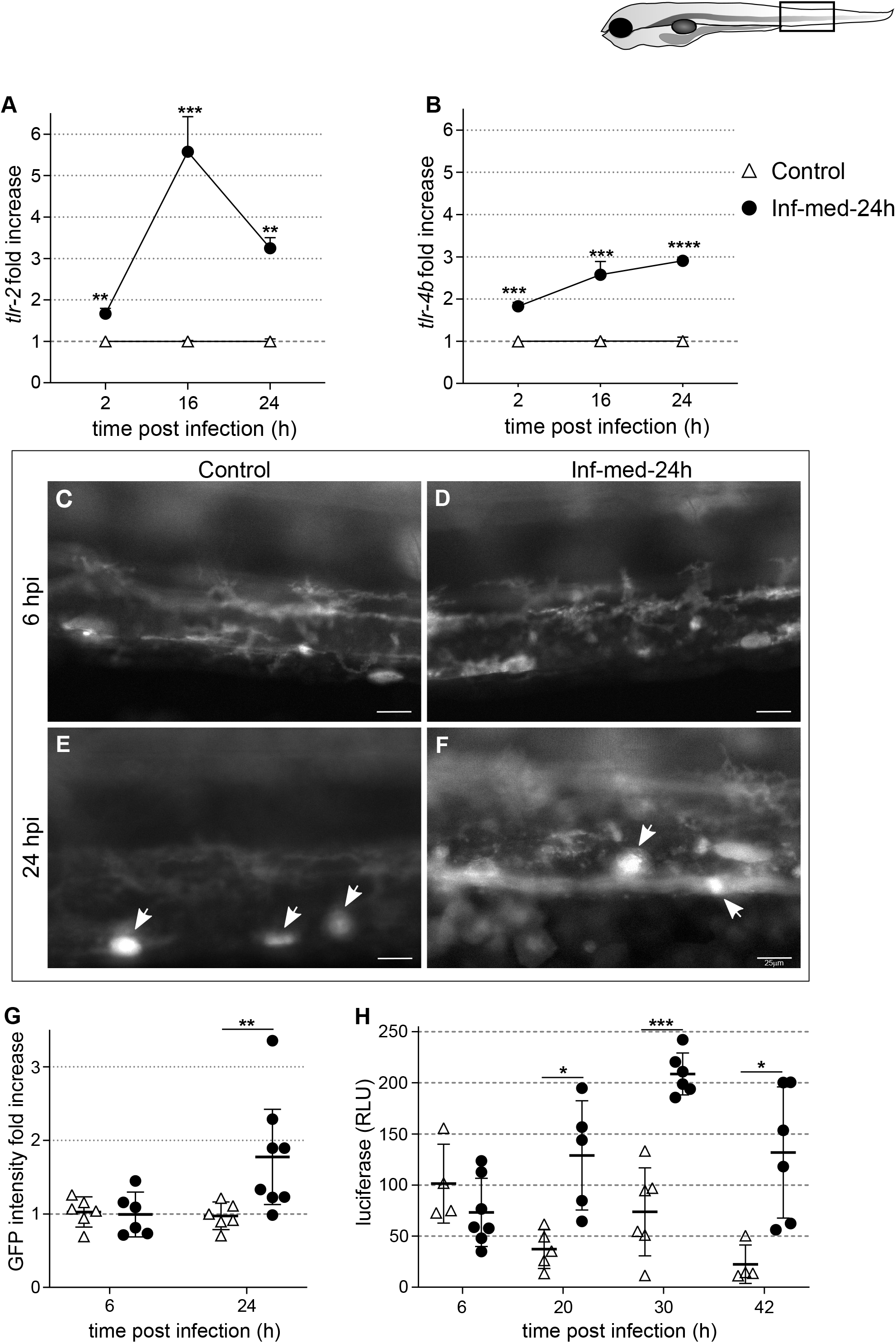
*S.epidermidis* infection upregulates *tlr-2* that activates NF-κB pathway. Transcriptional upregulation of *tlr-2* (**A**) and *tlr-4b* (**B**) was tested by real-time qPCR. (**A**) *tlr-2* showed significant upregulation in Inf-med-24h larvae as early as 2 hpi and the levels remained elevated till 24 hpi (**, p < 0.01; ***, p < 0.001). (**B**) *tlr-4b* mRNA expression was also upregulated upon infection. (***, p < 0.001). Data representative of three independent experiments. (**C** – **H**) *Tg*(*NFκB:GFP, Luc*) larvae were infected with Inf-med-24h and NF-κB activity was followed by GFP expression as well as luciferase activity. At 6 hpi, there was no discernible increase in NF-κB activity between the two experimental conditions (**C**& **D**). The intensity of GFP expression (**G**) and luciferase (**H**) activity was similar between the uninfected and infected larvae at this time-point. At 24 hpi, however, the infected larvae exhibited higher GFP expression than the control population (**E**vs **F**). (**G**) GFP intensities in similar ROIs were calculated and plotted as fold change in infected over uninfected larvae (**, p < 0.01) (n = 6 – 8 larvae). (**H**) The luciferase levels in infected larvae were significantly elevated 20 hpi onward (*, p < 0.05; ***, p < 0.001) (9-13 larvae). Box represents the area imaged. Images are representative of three independent experiments. Autofluorescent xanthophores are marked by white arrows. Quantification reflects combined data of three independent experiments. Scale bar, 25 μm.

Activation of the NF-kB pathway has been shown to occur as a consequence of activation of macrophages as well as upregulation of TLR signalling pathways (Dorrington and Fraser, 2019; Ivashkiv, 2011). We looked into the activation status of the NF-κB pathway in our model using the *Tg*(*NFκB:GFP, Luc*) transgenic reporter line, where an NF-κB responsive element drives the expression of GFP and luciferase. GFP intensities from similar ROIs of uninfected and Inf–med-24h larvae were imaged at different time-points post-infection (**Fig - 5C-F**). At 6 hpi, there was no discernible difference in GFP intensities between the control population and the infected population (**Fig - 5C-D**). However, at 24 hpi, Inf–med-24h larvae had significantly higher GFP signals compared to the controls, indicating activation of the NF-κB pathway (**Fig - 5E-F**). The intensities of GFP signal from these ROIs were further quantified and compared between infected and control population. At 6 hpi, there was no significant difference between the two populations. But at 24 hpi, the average GFP intensity of the ROIs from the infected population was ~1.8-fold higher than that of the control population (**, p = 0.008) (**Fig - 5G**). It must be noted here that the GFP intensities thus calculated are inclusive of the intensities contributed by autofluorescent xanthophores, which are similarly distributed in control and infected populations (as indicated by arrows). The presence of these auto-fluorescent cells therefore lead to under-representation of fold-increase brought about by NF-κB activation. We further corroborated these findings by quantitating the luciferase activity. As can be seen from **Fig - 5H**, there was no detectable increase in NF-κB levels at 6 hpi. By 20 hpi, Inf-med-24h larvae showed elevated luciferase levels, indicative of increased NF-κB activity (*, p = 0.046). Elevated NF-κB signalling was sustained in infected larvae till 42 hpi, as evinced by higher luciferase levels both at 30- and 42 hpi (*, p = 0.029, ***, p = 0.0008) (**Fig - 5H**).

### *S.epidermidis* infection provokes a pro-inflammatory host response

NF-κB is a cardinal mediator of inflammation. Given the elevated levels of NF-κB in our infection model, our next aim was to characterise the inflammatory profile of the larvae infected with *S.epidermidis*. Towards this objective, we analysed the mRNA levels of a select set of NF-κB responsive pro-inflammatory cytokines at different times post infection by real-time qPCR (**Fig - 6**). The mRNA levels of *il-1b* (**Fig - 6A**), *tnf-a* (**Fig - 6B**), *il-12b* (**Fig - 6C**) and *il-6* (**Fig - 6D**) were normalised against the house-keeping gene *b-actin*. As early as 2 hpi, mRNA levels of *il-1b* were significantly elevated (2.9-fold) in infected larvae. The *il-1b* levels in infected larvae peaked at 16 hpi, (16.5-fold rise) and then abated. At 24 hpi, the *il-1b* levels were still significantly high (6.5-fold) (***, p = 0.0003; ****, p = 4.9 × 10^−5^) (**Fig - 6A**). *tnf-a* mRNA levels showed a similar pattern – a rapid rise at 2 hpi (over 3-fold), with further increase at 16 hpi (over 11-fold) followed by slight abatement at 24 hpi (4-fold) (**Fig - 6B**) (*, p = 0.047; ***, p = 0.0008). We saw upregulation of *il-1b* and *tnf-a* mRNAs as early as 2 hpi (**Fig - 6A-B**), even though NF-κB activation was evident by 20 hpi (**Fig - 5H**). A likely reason for this apparent delay is the difference in the nature of the respective measurements. NF-κB activation was measured as function of mature reporter proteins (GFP and luciferase), whereas NF-κB responsive genes were detected at their mRNA levels. It is worth noting, that upregulation of the cytokines closely matches the increase in macrophage numbers (**Fig - 4**), indicating a precise and swift host immune response.

**Figure – 6:**
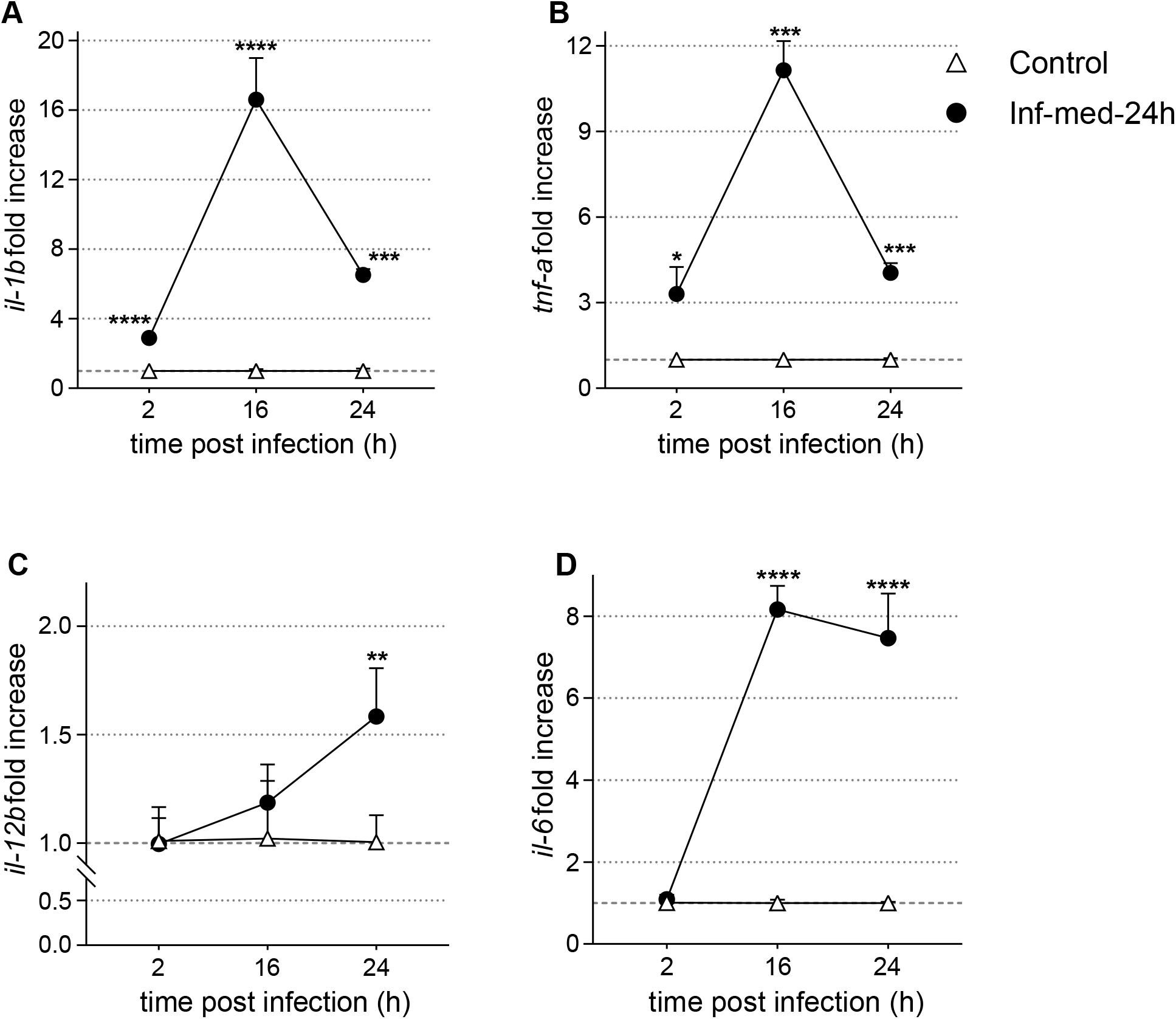
Zebrafish host response has a pro-inflammatory signature. mRNA levels of various pro-inflammatory cytokines were investigated in control and Inf-med-24h larvae by real-time qPCR. *il-1b* (**A**), *tnf-a* (**B**) and *il-6* (**D**) had similar expression profiles. All the three cytokines showed peak expression at 16 hpi. At 24 hpi, the levels of these cytokines had reduced but were still significantly elevated compared to uninfected controls. *il-12b* (**C**) levels between the two experimental groups were similar till 16 hpi. At 24 hpi, the infected larvae had a modest, but significant increase in *il-12b* levels. (*, p < 0.05, **, p < 0.01, ***, p < 0.001, ****, p < 0.0001). Representative of three independent experiments.

In contrast to *il-1b* and *tnf-a*, mRNA levels of both *il-12b* and *il-6* showed a delayed response in infected larvae (**Fig - 6C-D**). mRNA levels of *il-12b* remained unaffected till 16 hpi, and showed a mild, albeit significant elevation at 24 hpi (1.5-fold, **, p = 0.009) (**Fig - 6C**). The levels of *il-6* transcipts were similar to that of uninfected controls at 2 hpi. But by 16 hpi there was significant transcriptional upregulation of *il-6* transcription (8-fold increase) and it remained elevated till 24 hpi (7.5-fold increase) (****, p < 0.0001) (**Fig - 6D**).

### IL-6 signalling promotes resistance against *S.epidermidis* infection

IL-6 is a multifunctional cytokine which plays an important role in inflammation and immunity (Su et al., 2017). In order to elucidate the significance of the enhanced IL-6 signalling in zebrafish host response to *S.epidermis*, we sought to inhibit the IL-6 signalling pathway. Given that IL-6 signals through the JAK/STAT-3 pathway we inhibited the signalling with the help of two well-known STAT-3 inhibitors (WP-1066 and C-188-9) (Horiguchi et al., 2010; Jung et al., 2017). A range of concentrations were tested for both the inhibitors and WP-1066 was used for infection experiments at a final concentration of 12 μM (**Fig - 7A**). Prolonged exposure to C-188-9 resulted in significant larval mortality and hence this inhibitor was not pursued further (**Fig S2**). Uninfected larvae maintained in E3 with WP-1066 (inhibitor) or DMSO (vehicle control) showed no significant mortalities. In accordance to earlier observation, larvae exposed to low dose of bacteria for 6 hours (DMSO-Inf-low-6h) did not show any significant mortality compared to the uninfected controls (DMSO-Control). However, when IL-6 signalling was inhibited (Inh-Inf-low-6h), the same infection condition resulted in significant mortalities in larvae 54 hpi onward compared to their corresponding control population (Inh-Control) (*, p = 0.02). At 80 hpi, Inh-Inf-low-6h showed 65 % survival (**Fig - 7A**, closed squares) as opposed to 92 % and 98 % in DMSO-Inf-low-6h and Inh-Control respectively (****, p = 3 × 10^−6^) (**Fig - 7A**, open squares and closed triangles). The same trend was also observed in Inf-med-24h dose larvae. From 20 hpi, Inh-Inf-med-24h larvae showed significant mortality when compared to its uninfected counterpart (Inh-Control) (***, p = 9 × 10^-4^). At this time point, the survivals in the two infected populations were comparable (80 % DMSO-Inf-med-24h vs 79 % Inh-Inf-med-24h). However, by 54 hpi, Inh-Inf-med-24h group exhibited significantly lower survival values (51 %) compared to its untreated, DMSO-Inf-med-24h, counterparts (71 %) (***, p = 0.0008). At the end of the study, 57 % of larvae maintained in DMSO (DMSO-Inf-med-24h) survived as opposed to 25 % of Inh-Inf-med-24h (****, p = 1 × 10^−5^) (**Fig - 7A**, open and closed circles). This data clearly demonstrated that IL-6 signalling is an important component of the mechanism of resistance of zebrafish larvae against *S. epidermidis* infection.

**Figure – 7:**
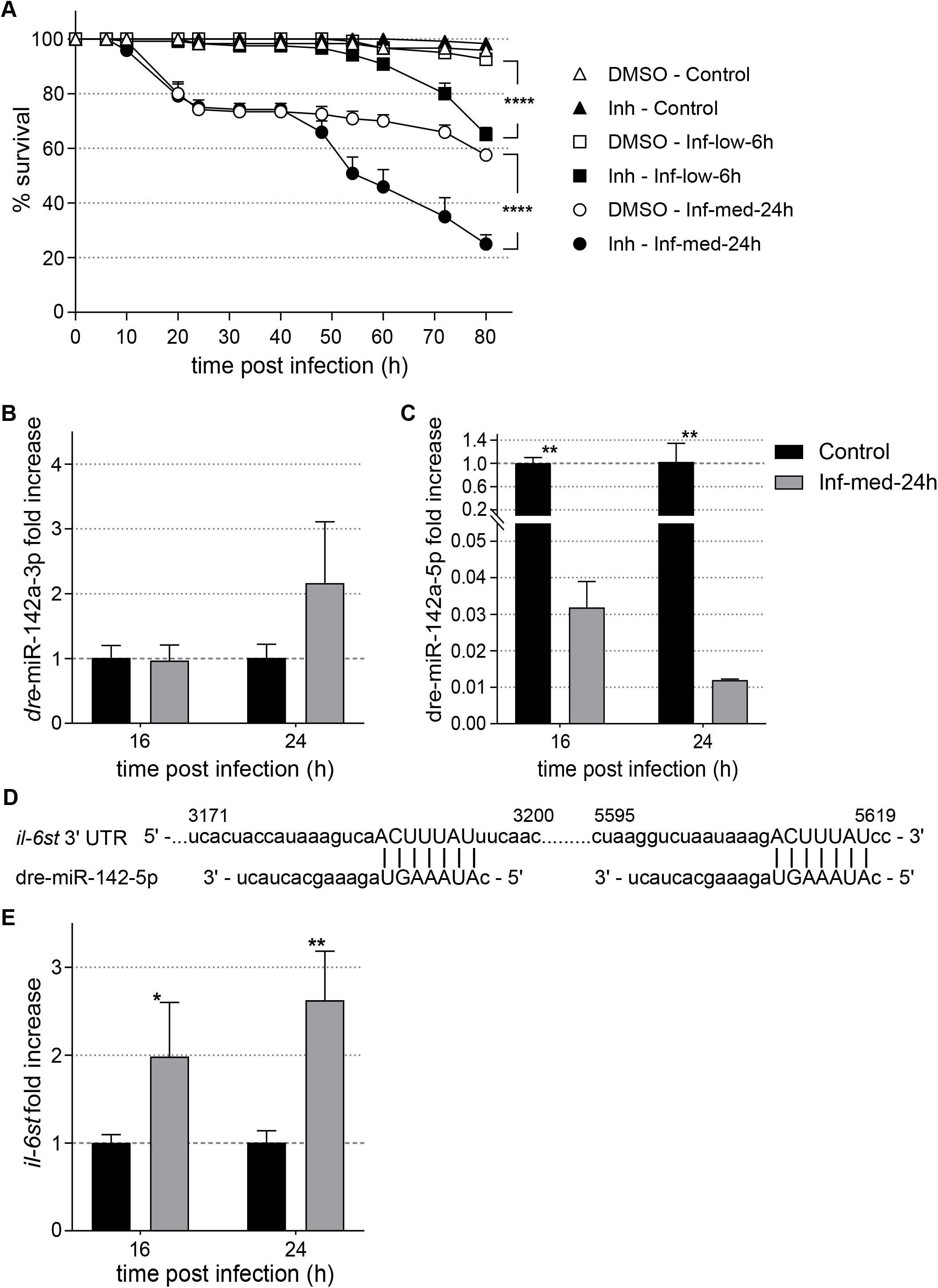
IL-6 signalling protects the host against infection and is regulated by *dre-miR-142-5p*. (**A**) Inhibition of IL-6 signalling rendered larvae more susceptible to *S.epidermidis* infection. Larvae (20/ well) infected with DMSO-Inf-low-6h were not affected by *S.epidermidis*. Inh-Inf-low-6h larvae, however, showed significantly higher mortalities. The same trend was observed when comparing mortality profiles of DMSO-Inf-med-24h and Inh-Inf-med-24h. (****, p < 0.0001). (**B** – **C**) qPCR showed that while *dre-miR-142-3p* (**B**) was unaffected by *S.epidermidis* infection, levels of *dre-miR142-5p* were drastically down-regulated (**C**). Inf-med-24h larvae showed significant down-regulation both at 16 hpi and 24 hpi (**, p < 0.01). (**D**) The 3’-UTR of *il-6st* is predicted to have two dre-miR-142a-5p binding sites: between 3188 – 3194 and 5611 – 5617. (**E**) Levels of *il-6st*, a target of dre-miR-142-5p, was significantly up-regulated in Inf-med-24h at 16- and 24 hpi (*, p < 0.05, **, p < 0.01). Representative of three independent experiments.

### miR-142-5p regulates IL-6 signalling by modulating IL-6 signal transducer

Given the importance of IL-6 signalling in the host immune response to *S.epidermidis* infection, we investigated the mechanism(s) of IL-6 regulation. The IL-6 signalling pathway is regulated by multiple miRNAs (Servais et al., 2019). miR-142 is one of the major miRNAs that has been shown to have important functions in infections and inflammation (Shrestha et al., 2017). Additionally, miR-142 has been shown to directly target various components of the IL-6 signalling pathway in humans and mouse (Liu et al., 2016; Sun et al., 2011; Wong et al., 2016). This miRNA has also been shown to impede clearance of *S.aureus* at wound sites, albeit by modulating small GTPases involved in neutrophil motility, and thereby altering neutrophil chemotaxis (Tanaka et al., 2017). Both the isoforms of miR-142 (dre-miR-142a-5p and dre-miR-142a-3p) are present in zebrafish and the latter has been shown to be necessary for neutrophil development (Fan et al., 2014). However, there is no evidence for interaction of these miRNAs with any gene of the IL-6 signalling pathway, in zebrafish. To investigate if dre-miR-142 was involved in the host response to *S.epidermidis* infection, we studied the miRNA levels of both dre-miR-142a-3p (**Fig - 7B**) and dre-miR-142a-5p (**Fig - 7C**). There was no significant change in the levels of dre-miR-142a-3p at both 16- and 24 hpi, suggesting that this miRNA was not involved in regulating *il-6* in this infection model. Levels of dre-miR-142a-5p, however, were significantly down-regulated in Inf–med-24h larvae. At 16 hpi, it was 0.032-fold (~ −30-fold) in infected larvae compared to controls, while at 24 hpi, the levels further decreased to 0.012-fold (~ −83-fold) (**, p = 0.002) (**Fig - 7C**). Given that there is no experimental or computational evidence of *il-6* being directly targeted by miR-142-5p, we looked at other genes of the IL-6 signalling pathway, namely IL-6 receptor alpha (IL-6Rα) and IL-6 signal transducer (IL-6st). IL-6 binds to IL-6Rα which triggers the recruitment of IL-6st that subsequently initiates the signal transduction (Su et al., 2017). While no predictions of a direct interaction was found between hsa-miR-142-5p and *il-6ra* in humans, a direct interaction between hsa-miR-142-5p and *il-6st* was predicted by the miRNA prediction software miRDB (Chen and Wang, 2020). Additionally, TargetScanFish (Ulitsky et al., 2012) predicted two dre-miR-142a-5p target sites in the zebrafish *il-6st* sequence (**Fig - 7D**). Hence, we decided to look at the mRNA levels of *il-6st* in this infection model (**Fig - 7E**). The *il-6st* levels of Inf-med-24h larvae, showed a very tight inverse correlation with dre-miR-142a-5p levels at both the time-points we checked. Compared to uninfected controls, at 16 hpi *il-6st* showed ~1.9-fold elevation (*, p = 0.044), that increased to ~2.6-fold by 24 hpi (**, p = 0.005). While in vitro assay showing direct binding between dre-142a-5p and *il-6st* is required for confirmation, our data strongly suggests that the upregulation of IL-6 signalling pathway in this infection model involves dre-miR-142a-5p. Under uninfected condition, dre-miR-142a-5p likely interacts with, and down-regulates *il-6st*, keeping the pathway under check. *S.epidermidis* infection causes severe down-regulation of dre-miR-142a-5p, thus enabling upregulation of the IL-6 signalling pathway.

## Discussion

*S.epidermidis* has recently emerged as an important opportunistic pathogen that causes frequent nosocomial infections. Owing to the natural occurrence of the bacteria as commensals on human skin, it is a common cause of acute sepsis in premature neonates and immunocompromised individuals (Sabaté Brescó, Marina et al., 2017). It is the single largest cause of infections in implants and indwelling medical devices (Otto, 2009; Uçkay et al., 2009). Various implant models have been established to study infection as well as clearance of this bacterium (Sabaté Brescó, M. et al., 2017; Walker et al., 2019). While these studies have demonstrated the conditions that promote infections, host immune responses to *S.epidermidis* have remained less well studied.

Zebrafish has emerged as an effective model for studying host-pathogen infections in vivo (Gomes and Mostowy, 2020; Renshaw and Trede, 2012) and zebrafish models of *S.aureus* infection have shed light on various virulence mechanisms (Prajsnar et al., 2008; Prajsnar et al., 2012). Veneman *et al.*, have recently established a high-throughput screening system where zebrafish embryos were injected with *S.epidermidis* (Veneman et al., 2013). Here, we have developed a novel *S.epidermidis* bath immersion systemic infection model for zebrafish to elucidate the host immune responses. *S.epidermidis* recognition and bacterial clearance has been shown to be mediated by TLR-2 in a mouse model (Strunk et al., 2010). The zebrafish host response also showed elevation of *tlr-2* transcription and NF-κB activation indicating that Tlr-2 signalling was upregulated. There was marked inflammation with upregulation of various pro-inflammatory cytokines such as *il-1b*, *tnf-a* and *il-6*. The infected larvae exhibited a rapid cellular response that was driven by macrophages. There was over 2-fold increase in macrophage numbers within the first two hours of infection. Macrophage numbers remained elevated throughout the infection period and beyond indicating a role in clearance of infection. We further show the key role of IL-6 signalling in modulating inflammation and thus infection prognosis, and discover miR-142-5p as a potential candidate that can likely regulate this pathway.

The new non-invasive bath immersion model that we have established is experimentally highly amenable. As expected, compared to experiments that inject the bacterium into the embryo (Veneman et al., 2013), relatively higher inoculum was required to obtain reproducible infection phenotypes in this model. For most of our experiments we used a dose that induced marked phenotypes of infection and host immune responses (Inf-med-24h) in order to facilitate the analyses. However, it is worth noting that the even the lowest dose (Inf-low-6h), that caused little morbidity and no mortality resulted in bacterial colonisation with a concomitant measurable host immune response indicated by significant leukocytosis. These suggest that *S.epidermidis* does infect the larvae at lower doses and a robust host immune response efficiently clears the infection. Hence this model is usable at lower MOIs for experiments that do not require overt morbidity and mortality phenotypes. The mechanisms of host immune response to non-pathogenic low doses could potentially differ from the inflammatory mechanisms that we have uncovered in our pathogenic model and may be more relevant to explaining the host-pathogen interface in asymptomatic commensal relationships. This aspect would be interesting to explore in further studies.

Twenty-four-hour long exposure to bacterial broths does not result in higher colonisation, in either of the infection doses (Inf-low-24h and Inf-med-24h). This indicates that after the initial colonisation, rapid immune responses can efficiently bulwark against further accretion of infections. Macrophages have been shown to play an important role in combatting *S.aureus* infection in murine, human as well as zebrafish hosts (Kubica et al., 2008; Prajsnar et al., 2008; Verdrengh and Tarkowski, 2000). Similar to *S.aureus* infection, *S.epidermidis* infection in zebrafish resulted in the involvement of myeloid populations, predominantly the macrophages. Within 2 hours post infection we were able to measure significant increase in the number of macrophages in both the doses we analysed. In our experiments, neutrophils do not appear to play a major role in the innate response and macrophages act as the primary innate responder cells. The initial cellular response to the presence of *S.epidermidis* is found to be comparable in magnitude across all the doses that we have tried, a 2-fold increase in macrophage numbers in 2 hours. This, perhaps, is the primary rapid response. The further gradual increase observed occurs commensurate with increasing bacterial exposure in dose and time. This increase, provoked by prolonged bacterial colonisation of the host, probably has an important role in clearance and repair. This function is also suggested by the nearly 16 hours delay in return of macrophage numbers to control levels beyond the bacterial clearance in the fish infected with Inf-low-6h. *S.epidermidis* strains that are unable to form biofilm are more susceptible to phagocytosis (Shiro et al., 1994), whereas *S.aureus* and *S.epidermidis* biofilms can inhibit the phagocytic activity of macrophages (Spiliopoulou et al., 2012; Thurlow et al., 2011). Given that we see an increase in macrophage numbers while systemic bacterial counts are decreasing, it would be interesting to see if the bacteria on prolonged colonisation activate any of the biofilm pathways.

Macrophage phagosomes recruit Tlr-2 that gets activated by Gram-positive but not Gram-negative bacteria (Underhill et al., 1999). Tlr-2 is essential for neutrophil and macrophage recruitment post-*S.aureus* infection (Fournier, 2012). In the context of *S.epidermidis* infection, Tlr-2 has been shown to be involved in recognition as well as clearance of the bacteria (Strunk et al., 2010). *S.epidermidis* infection in this model recapitulated those findings by showing an upregulation of *tlr-2*. We also observed moderate elevation of *tlr-4b* levels. Although the role of *tlr-4* in *S.epidermidis* infection is not known, in *S.aureus* infection of murine brain while *tlr-2* is central for activating downstream immune responses, *tlr-4* is also important for leukocyte apoptosis and cytokine production (Stenzel et al., 2008). Degradation of *S.aureus* within phagosomes can often times result in activation of Tlr-8 signalling in the endosomal compartment. In such a scenario, IRF5 pathway is activated that eventually results in induction of IFN-β, a classical antiviral cytokine. Tlr-2 antagonises Tlr-8-IRF5 signalling axis (Bergstrøm et al., 2015). Given that *tlr-2* was elevated and the levels of *tlr-8a* and *tlr-8b* were unaffected in the infected larvae, it is likely that *S.epidermidis* infection did not activate the Tlr-8-IRF5 signalling cascade.

Convergence of Tlr signalling into NF-κB activation is well documented (Kawai and Akira, 2007). *S.epidermidis* lysates have been shown to activate NF-κB signalling in human and mouse epithelial cells (Val et al., 2015). Staphylococcal infections lead to pro-inflammatory immune response in a myriad of host models (Askarian et al., 2018; Fournier and Philpott, 2005). In our model, all the NF-κB responsive pro-inflammatory cytokines that we tested were elevated. The mRNA levels of *il-1b*, *tnf-a* and *il-6* showed stark increase, while *il-12b* showed a mild but significant elevation. *S.aureus* infection has been shown to induce *il-1b* in murine brain abscess models (Cho et al., 2012; Kielian et al., 2004). Spiliopoulou *et al.*, showed that human cell-lines infected with *S.epidermidis* induced IL-1β (Spiliopoulou et al., 2012), and we report elevation of the same in an in vivo model. They further showed that induction of IL-1β, TNF-α and IL-6 were unaffected by biofilm formation; but IL-12 elevation was dampened when *S.epidermidis* formed biofilms (Spiliopoulou et al., 2012). In light of this observation, it will be interesting to investigate whether the profile of *S.epidermidis* infection in our model resembles biofilm or planktonic cultures. TNF-α is another cytokine that is upregulated upon *S.aureus* infection in mouse models (Kielian et al., 2004; Yimin et al., 2013). *S.epidermidis* infection also induces this cytokine in human cell-culture (Spiliopoulou et al., 2012) and mouse and zebrafish models (Strunk et al., 2010; Veneman et al., 2013). Our bath immersion model recapitulates this feature of the infection. Tlr-2 activation by *S.aureus* and *S.epidermidis* infections result in upregulation of IL-6 levels in mouse (Strunk et al., 2010; Yimin et al., 2013). Neonatal sepsis due to *S.epidermidis* infection show elevated levels of IL-6 (Tröger et al., 2018). In zebrafish, *S.epidermidis* infection resulted in heightened *il-6* levels.

IL-6 is a well-known pleiotropic cytokine, with pro-as well as anti-inflammatory effects (Su et al., 2017). Hence we sought to look at the significance of elevated *il-6* levels in this infection model. Evidences suggest that the IL-6 classic signalling leads to the regenerative and anti-inflammatory effects of IL-6, while IL-6 trans-signalling is responsible for pro-inflammatory phenotypes (Chalaris et al., 2011). Tlr-2 activation has been shown to induce IL-6 trans-signalling in monocytes (Flynn et al., 2019). IL-6 trans-signalling has also been implicated in mediating inflammatory responses in various models of infection and endotoxin challenge (Hunter and Jones, 2015). While IL-6 signalling promoted a Tlr-4 dependent pro-inflammatory response in a murine endotoxin model (Greenhill et al., 2011), IL-6 signalling in *Toxoplasma gondii* had an anti-inflammatory effect (Silver et al., 2011); both of which were detrimental to animal survival. In our model however, inhibition of IL-6 signalling using STAT-3 inhibitor, rendered the zebrafish more susceptible to *S.epidermidis* infection induced mortality. This is reminiscent of *S.aureus* infection reports which showed that elevated IL-6 signalling led to higher survival of mice (Onogawa, 2005) and lack of IL-6 signalling resulted in greater infection loads (Hume et al., 2006). Onogawa *et al*. showed that enhancement of IL-6 signalling led to augmentation of various cytokines such as IL-1α, IL-4 and TNF-α. While upregulation of IL-6 during *S.epidermidis* infection has been shown ex vivo and in vivo, to the best of our knowledge, this is the first report of a protective role of IL-6 signalling in this infection. It will be interesting to investigate whether the protective effect of IL-6 signalling in *S.epidermidis* infection gives rise to a similar profile of immune modulators as that seen in *S.aurues* infection.

Given the significance of IL-6 signalling in this model, we sought to explore the possible mechanism(s) of its regulation. miRNA-mediated modulation is one of the important mechanisms of regulation of different genes of this pathway (Tanaka et al., 2016). miR-142 is one of the several miRNAs that has been shown to be involved in the IL-6 signalling pathway (Servais et al., 2019). miR-142-3p has been reported to target *il-6* in models of endotoxin-challenge and age-related studies (Liu et al., 2016; Sun et al., 2011). Regulation of *il-6* by miR-142-3p in an infection setting has not been previously reported. However, miR-142-3p has been implicated in affecting phagocytosis upon *S.aureus* as well as mycobacterial infections (Bettencourt et al., 2013; Tanaka et al., 2017; Xu et al., 2013). In our model dre-miR-142a-3p levels remained unaffected, indicating this miRNA is unlikely to be involved in regulating the immune response to *S.epidermidis* infection. The role of miR-142-5p, in contrast, has remained relatively obscure. It has been shown to directly target *il-6st* in THP-1cells under conditions of temperature stress (Wong et al., 2016) and during adaptive hypertrophy in mice (Sharma et al., 2012). Nothing has been shown about the role of this miRNA or the role of its interaction with *il-6st* in infections or in immune responses. Here, we show for the first time a likely involvement of miR-142-5p in regulating IL-6 signalling pathway upon an infection. We see an inverse relationship between levels of dre-miR-142a-5p and of *il-6st*. Thus, it is likely that in zebrafish also dre-miR-142a-5p regulates *il-6st* by directly targeting it. miRNA profiling in bovine models gave conflicting reports regarding the effect of *S.aureus* infection on bta-miR-142-5p expression (Jin et al., 2014; Luoreng et al., 2018; Sun et al., 2015). Here we show drastic down-regulation of dre-miR-142a-5p with concomitant upregulation of its target gene *il-6st*. Thus we propose that the host response to *S.epidermidis* infection is characterised by a NF-κB dependent pro-inflammatory response and an upregulation of IL-6 signalling with the likely involvement of dre-miR-142a-5p.

In conclusion, we have established a novel bath immersion model of *S.epidermidis* in zebrafish and have elucidated several key features of the host immune response. The model is versatile in that it allows experimentation with low as well as high MOIs and can serve as a platform to address various aspects of host pathogen interactions and drug screens. *S.epidermidis* infection results in leukocytosis along with upregulation of Tlr-2 signalling. The infection causes a strong pro-inflammatory response and sustained upregulation of IL-6 signalling pathway that has a protective effect. We also identify a hitherto undescribed immune function of dre-miR-142-5p and its interaction with *il-6st* which might be a key factor in this regulation.

## Material and Methods

### Zebrafish reporter lines and maintenance

Zebrafish were maintained at 28 °C with a 14-h light/ 10-h dark cycle. Embryos were collected by natural spawning and maintained at 28.5 °C in E3 medium. For imaging experiments, phenylthiourea (0.003 %) was added to 1-dpf embryos in order to avoid melanisation. Wild-type (Tü) strain and three transgenic reporters were used: *Tg*(*8xHs.NF*κ*B:GFP,Luciferase*) (hereafter known as, *Tg*(*NFκB:GFP, Luc*)), *Tg(pu.1:Gal4.UAS RFP)* (subsequently referred to as *Tg*(*pu.1:RFP*)) and *Tg(mpeg1:Kaede)* in this study. All the experimental procedures have been approved by the Institutional Biosafety and Animal Ethics committees.

### Bacterial strain

*Staphylococcus epidermidis* (strain O-47) (Heilmann et al., 1996) was grown in TSB supplemented with 1% NaCl. It was streaked onto HiCrome Staph Selective Agar (Himedia) as well as TSA plates. The *S.epidermidis* colonies were blue in colour in the Staph Selective media.

### Establishment of infection

An overnight culture of *S.epidermidis* was grown and 1: 50 inoculum made on the day of infection. The bacteria were grown at 28.5 °C till O.D_600_ 1.0 and pelleted down by centrifugation at 4000 rpm for 10 min. The pellet was resuspended in sterile PBS and washed twice, and resuspended in E3 medium. In experiments using heat-killed bacteria, the final *S.epidermidis* suspension was incubated at 95 °C for 5 min.

Zebrafish larvae (4 dpf) were infected by exposure to *S.epidermidis* in bath immersion. Each well of a 24-well plate had 15 – 20 larvae in 2 ml E3 medium. The zebrafish larvae were exposed to varying concentrations (5 x 10^8^ CFU/ ml, 1 × 10^9^ CFU/ ml and 1 × 10^10^ CFU/ ml) of *S.epidermidis* for either 6 h or 24 h. Subsequently, the larvae were removed from infected E3 media, washed twice in E3 media and transferred to fresh media for further study. The infection dose was validated by CFU plating. The time of addition of the bacterial inoculum was considered as time zero and time-points were calculated as hours post-infection thereafter. For survival studies, triplicate wells of each infection dose were monitored till 80 hpi.

For inhibition of IL-6 signalling, two STAT-3 inhibitors (STAT-3 inhibitor III (WP-1066) and STAT-3 inhibitor XIII (C-188-9) were tested (Merck, Sigma) and WP-1066 was used for infection studies at a final concentration of 12 μM. WP-1066 was added to E3 medium 30 min prior to infection and the larvae were maintained in that medium during the entire period of the mortality study. Equivalent volume of DMSO was added to wells with larvae serving as vehicle control.

### CFU counts

From each experimental condition, 3 larvae were randomly selected at different time points post-infection. Individual larva was anesthetized with a lethal dose of Tricaine, homogenised mechanically in 1 % Triton-X100 in PBS and multiple dilutions of the lysate plated in triplicate. The plates were incubated at 37 °C for 16-20 h, colonies counted and CFU calculated accordingly.

### Quantitative PCR analysis

Total RNA was extracted from a pool of 10 – 15 embryos at different time points from each experimental condition using Trizol (Thermo Fisher Scientific) following the manufacturer’s protocol. cDNA was prepared by reverse transcription using SuperScript Vilo (Thermo Fisher Scientific). Gene-specific primers were designed using Universal Probe Library (Roche) and real-time PCR performed using GoTaq qPCR Master mix (Promega) on ABI QuantStudio 7 Flex Real Time PCR System. *b-actin* was used as a housekeeping gene for normalisation. For miRNA qPCR, cDNA was prepared using miRNA 1-st strand cDNA synthesis kit (Agilent). Mature miRNA sequence and Universal reverse primer (Agilent) was used as forward and reverse primers respectively. The Tm of the qPCR reaction was adjusted to 59 °C (Busk, 2014). dre-miR-451 was used for normalisation. The primer sequences are tabulated in **Supplementary Table - S3**.

### Imaging

Larvae were anaesthetized with 0.01% tricaine, embedded in 1.2 % low-melting-point agarose, and imaged using Nikon TiEB (20x) and Leica DMi8 (20x). A total of 20 – 30 z-stacks of 2.5 μm thickness was captured. Image analysis was done using Fiji software (Schindelin et al., 2012). NFκB activity of infected and uninfected *Tg*(*NFκB:GFP, Luc*) larvae was visualised by imaging similar regions of uninfected controls and infected larvae at different time points post-infection. For quantification of GFP intensity similar region of interest (ROI) was selected from each larva and intensity per area calculated. The final signal intensities for were determined following background subtraction. The intensities were normalised to control values and plotted as fold increase (n = 6 - 8 larvae).

In order to enumerate leukocytes, multiple fields of uninfected and infected *Tg*(*pu.1*:*RFP*) larvae, spanning the trunk and tail regions, were imaged at different time-points post-infection. The total number of RFP-positive cells in similar fields of uninfected and infected larvae were counted manually and an average number of leukocytes in that particular field was calculated. Double-positive larvae, obtained from mating *Tg*(*pu.1*:*RFP*) × *Tg*(*mpeg1*:*Kaede*), was used to count number of macrophages (RFP and Kaede (green) positive) and neutrophils (RFP positive) (n = 9 - 12 larvae).

### Luciferase measurements

*Tg*(*NFκB:GFP, Luc*) larvae were used to quantify NFκB activity by assaying luciferase activity as reported (Kuri et al., 2017). Briefly, individual larva were transferred to each well of a 96-well optical bottom plate (Nunc) in 50 μl of E3 medium, without methylene blue (n = 9 – 13 larvae). The medium was supplemented with 1 mM beetle luciferin potassium salt solution (Promega) and bioluminescence assayed at room temperature using Varioskan Flash (Thermo Fischer).

### Statistical Analysis

Data were plotted using GraphPad PRISM. Multiple t-tests and two-way ANOVA was performed for statistical analysis. p < 0.05 at a confidence interval of 95% was considered significant.

## Author Contributions

S.B designed the project, analysed the data and wrote the manuscript. D.P.T. carried out most of the experiments and analysed the data. P.K.K. assisted D.P.T in experimentation and data analysis. R.K.S. collaborated on the project.

## Acknowledgements

We thank Maria Leptin, Francesca Peri and Graham Lieschke for their generous gifts of *Tg*(*NFκB:GFP, Luc*)), *Tg*(*pu.1:RFP*) and *Tg*(*mpeg1*:*Kaede*) reporter lines respectively; Harapriya Mohapatra and Martina Rembold for critical comments on the manuscript; Ramanujam Srinivasan and Renjith Mathew for insightful discussions throughout the project; and School of Biological Sciences, NISER for departmental support. The work was supported by Ramanujan Fellowship (SB/S2/RJN-174/2014) and DST-ECRA (2017-000528) grant to S.B; intramural funds from Institute of Life Sciences (an autonomous institute of DBT) and SERB (EMR/2016/003780) grant to R.K.S.

