## Supplementary files S1-S3 for "IL-6 signalling protects zebrafish larvae during *Staphylococcus epidermidis* infection in a novel bath immersion model"

#### Figure - S1

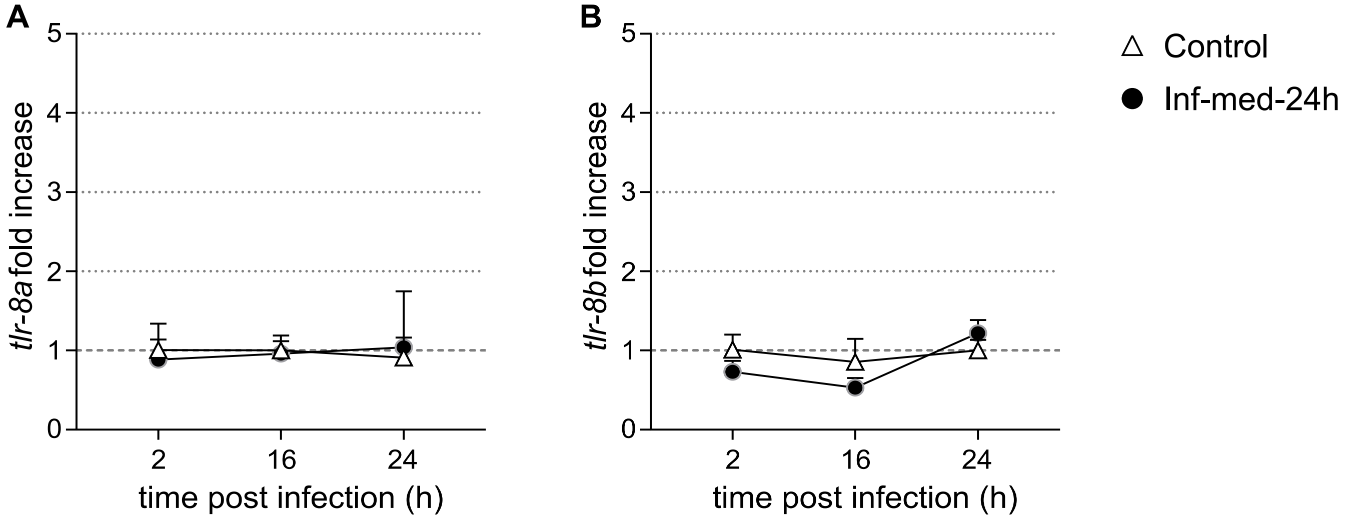


**Figure – S1**: *tlr-8a and tlr-8b are unaffected by S.epidermidis infection*. Transcriptional upregulation of *tlr-8a* (**A**) and *tlr-8b* (**B**) was tested by real-time qPCR. There was no significant difference in *tlr-8a* and *tlr-8b* levels between uninfected larvae and those infected with Inf-med-24h.

### Figure - S2

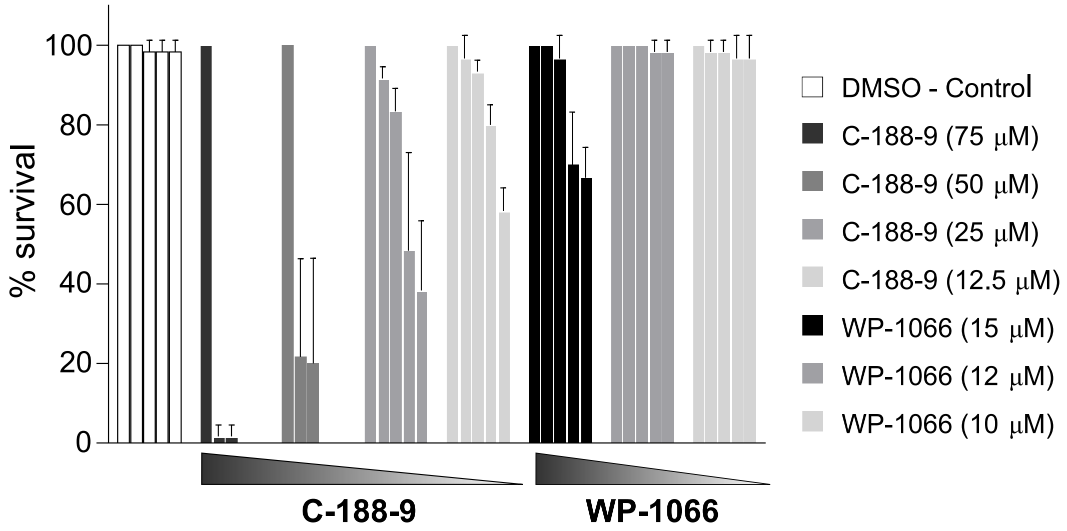


**Figure – S2**: Effect of Stat-3 inhibitors C-188-9 and WP-1066 on larval mortality. Larvae, 4 dpf, were maintained in varying concentrations of C-188-9 and WP-1066 and mortality observed at 0 h, 24 h, 48 h, 72 h and 80h. Compared to larvae maintained in DMSO, those maintained in C-188-9 showed significant mortality, even at the lowest dose (12.5 μM) tested (****, $p < 0.0001$). WP-1066 was less toxic to zebrafish larvae. While the highest concentration of WP-1066 (15 μM) caused significant mortality as compared to DMSO controls (****, $p < 0.0001$), the lower doses (12 μM and 10 μM) showed no significant death.

**Supplementary Table – S3.** Sequence of primers used for qPCR

| **Gene name** | **Primer sequence (5' – 3')** |
|---|---|
| *b-actin* | Forward: GATCTTCACTCCCCTTGTTCA<br>Reverse: GGCAGCGATTTCCTCATC |
| *il-1b* | Forward: TGAAGTCACCATAGCTCCAAAAA<br>Reverse: GCATGTCGCATCTGTAGCTC |
| *tnf-a* | Forward: AGGCAATTTCACTTCCAAGG<br>Reverse: AGGTCTTTGATTCAGAGTTGTATCC |
| *il-6* | Forward: AAGGGGTCAGGATCAGCAC<br>Reverse: GCTGTAGATTCGCGTTAGACATC |
| *il-6st* | Forward: TTCATTCCAAGATCAAATGACG<br>Reverse: TCCATCATGAACCCACCAC |
| *il-12b* | Forward: CGCTGTAGGAAACGCAAAA<br>Reverse: GGAGACTTTGTGTGCGGTAAG |
| *tlr-2* | Forward: AGACACAATAATGGCAGTCAGG<br>Reverse: TACATGTCTGGGAGCACTCG |
| *tlr-4ba* | Forward: CATGCATGGGAAGAAACCTC<br>Reverse: CCAGAGTTTGAACCGAGGAA |
| *tlr-8a* | Forward: TGCCTTCATCACCTACGACA<br>Reverse: GATCGGGAGGAAGAGTTCG |
| *tlr-8b* | Forward: CGAGTGTCTTGCAACTGGAC<br>Reverse: TTGAGGCAAGTGATGTTCTCC |
| dre-miR-451a | Forward: CCGTTACCATTACTGAGTT |
| dre-miR-142-5p | Forward: ACATAAAGTAGAAAGCACTACT |
| dre-miR-142-3p | Forward: GTAGTGTTTCCTACTTTATGGA |
